# *Toxoplasma gondii*-induced NETs are mediated by CR3 engagement and the vital non-lytic NETosis pathway

**DOI:** 10.64898/2026.11.25.769188

**Authors:** Marco A. Martinez, Jaden Ernest S. Valle, Tatiane S. Lima

## Abstract

Approximately one-third of the global human population harbors an infection with the protozoan parasite *Toxoplasma gondii*. The immune system exploits neutrophils and one of their effector mechanisms, neutrophil extracellular traps (NETs), as an attempt to capture and neutralize *T. gondii*. NETs, composed of DNA, myeloperoxidase, citrullinated histone 3, and elastase, play a dual role in pathogen entrapment and amplification of the immune response. Traditionally, NET release was considered to result in neutrophil cell death. However, studies have identified an alternative process, vital NETosis, that allows NET formation while maintaining cell viability. Although *T. gondii* induces NET formation, the associated molecular mechanisms are not fully understood, and the neutrophil fate is unknown. In this study, the promyelocytic cell line HL-60 was differentiated into neutrophil-like cells (NLCs) and infected with type I (RH) or type II (PA7) *T. gondii*. Extracellular DNA was found in increasing concentrations over time in both type 1 and type 2 *T. gondii*-infected NLCs. Immunofluorescence staining for DNA and citrullinated histone 3 at 4, 6, and 16 hours post-infection confirmed NET formation by confocal microscopy. Cell viability at 16 hours post-infection was assessed by live/dead staining and extracellular ATP measurement, revealing that most NET-releasing neutrophils retained membrane integrity, consistent with vital NETosis. Additionally, immunofluorescence microscopy at 15 and 30 minutes further confirmed the fast release of nuclear DNA, a hallmark of vital NETosis. The use of opsonin-containing human serum and an anti-CD11b blocking antibody identified complement receptor 3 (CR3) as a receptor mediating *T. gondii*-induced vital NETosis. These findings demonstrate that *T. gondii* triggers vital NETosis in neutrophils through a CR3-dependent mechanism, advancing our understanding of the innate immune response to this globally prevalent parasite and laying the groundwork for future investigations into the molecular pathways that may be therapeutically targeted to support neutrophil-mediated immunity during infection.

## Introduction

Approximately one-third of the world’s population is infected with the obligate intracellular protozoan *Toxoplasma gondii*, the causative agent of toxoplasmosis (Abi Abdallah et al., 2012; Bliss et al., 2001; Kochanowsky & Koshy, 2018). In immunocompetent individuals, infection is typically asymptomatic, though some may develop fever, swollen lymph nodes, or other flu-like symptoms (Kochanowsky & Koshy, 2018). A competent immune response induces a critical stage conversion in the parasite, shifting it from the aggressively replicating tachyzoite form to the less virulent bradyzoite form, which persists in tissue cysts (Bliss et al., 2001; Halonen & Weiss, 2013; Mukhopadhyay et al., 2020). In immunocompromised individuals, however, this protective stage conversion fails, allowing the parasite to reactivate and replicate unchecked (Biswas et al., 2017; Zhao et al., 2014; Bergersen et al., 2024). This reactivation can produce severe, potentially fatal conditions, including toxoplasma encephalitis, myocarditis, and pneumonitis. Tachyzoites can also infect placental cells and breach the placental barrier, placing pregnant women at risk of fetal infection and miscarriage (Rudzki et al., 2021).

The immune mechanisms mediating stage conversion center on the production of interferon-gamma (IFN-γ) by NK cells and T lymphocytes, which, in turn, depends on interleukin-12 (IL-12) produced by dendritic cells, macrophages, monocytes, and neutrophils (Bliss, 1999; Bliss et al., 2001; Miranda et al., 2021). Neutrophils are among the first cells recruited to the site of *T. gondii* infection in the small intestine, where they directly eliminate the parasite through both oxygen-dependent and oxygen-independent mechanisms and contribute to mucosal immunity (Bergersen et al., 2023, 2024; Bliss et al., 2001; Lima et al., 2018). Consistent with this, neutrophil depletion prior to infection in mouse models results in higher parasite burden, attributable to weakened type 1 immune responses characterized by reduced IFN-γ and IL-12 production (Biswas et al., 2017; Bliss et al., 2001). The innate immune system relies on complement proteins and antibodies to opsonize pathogens and facilitate their destruction by phagocytes (Janeway et al., 2001.; Segal, 2005). The complement system relies heavily on C3b and iC3b as the main effector molecules to bind pathogens and mark them for destruction, while C3b also activates the complement system further downstream (Sikorski et al., 2021). Phagocytes, such as neutrophils, detect these complement proteins through a class of receptors known as complement receptors (Lamers et al., 2021). C3b is detected by complement receptor 1 (CR1) and complement receptor 3 (CR3), while iC3b is detected by CR3 and complement receptor 4 (CR4) in neutrophils (Gordon et al., 1987). CR3 is the most highly expressed complement receptor in human neutrophils (Erdei et al., 2019).

In addition to cytokine secretion and direct killing, neutrophils control *T. gondii* in part by forming neutrophil extracellular traps (NETs), web-like structures composed of DNA decorated with histones, elastase, myeloperoxidase (MPO), and other antimicrobial molecules (Abi Abdallah et al., 2012; Bliss et al., 2001; Lima et al., 2018). *T. gondii*-induced NETs have been shown to promote the systemic release of cytokines, including IFN-γ, TNF-α, IL-17, IL-12, and IL-10, from CD4+ cells and peripheral blood mononuclear cells, as well as to increase plasma levels of the chemokines CXCL8 and CCL4 (Miranda et al., 2021). These studies highlight a more extensive immunomodulatory role for NETs in coordinating both innate and adaptive responses to infection.

NETs were first described in 2004 and initially understood as a form of programmed cell death, termed NETosis, in which neutrophils expel nuclear DNA into the extracellular space as a terminal antimicrobial act (Brinkmann et al., 2004; Jorch & Kubes, 2017; Kubes, 2018; Liew & Kubes, 2019; Yipp & Kubes, 2013). This understanding was refined when a study examining neutrophil responses to *Staphylococcus aureus* demonstrated that NETs could be released without plasma membrane disruption or cell lysis (Pilsczek et al., 2010; Yipp et al., 2012). In this mechanism, termed vital NETosis, the nuclear envelope separates, and vesicles containing nuclear DNA are extruded into the extracellular space, where they rupture to release packed chromatin and antimicrobial molecules, while the neutrophil remains viable and retains effector functions such as chemotaxis and phagocytosis (Pilsczek et al., 2010a; Yipp & Kubes, 2013). By contrast, suicidal NETosis proceeds through superoxide formation, which leads to plasma membrane rupture and cellular death, eliminating these downstream functions (Remijsen et al., 2011). NET formation is traditionally confirmed experimentally by detecting citrullinated histone 3 (Cit-H3) in the extracellular space, as histone citrullination is required for chromatin decondensation during NET release (Chen et al., 2021;; Li et al., 2010). In vitro studies of NET formation have used HL-60 promyeloblasts, which can be differentiated into neutrophil-like cells (NLCs) (Bergersen et al., 2023; Brinkmann et al., 2004; Tasseff et al., 2017). Supplementation of differentiation media with retinoic acid enhances both differentiation and NET production (Tasseff et al., 2017).

*T. gondii* replication rates, host immune responses, and parasite immune evasion strategies vary significantly across clonal lineages, thus highlighting the importance of investigating multiple lineages in experimental work. In North America and Europe, the predominant lineages are types I and II, with type II responsible for most human and livestock infections (Kochanowsky & Koshy, 2018; Zhao et al., 2014). Although coculture of healthy donor neutrophils with *T. gondii* tachyzoites has been shown to induce NET formation (Abi Abdallah et al., 2012; Bliss et al., 2001), the cellular and molecular mechanisms that mediate this response remain not fully understood. Here, we report that *T. gondii*-induced NETosis proceeds through a fast CR3-mediated vital NETosis pathway. The characterization of this mechanism deepens our understanding of how neutrophils remain functionally active after nuclear DNA extrusion and may clarify the contribution of NETs to the early cytokine environment during acute *T. gondii* infection.

## Material and Methods

### Neutrophil-like cell differentiation

The promyeloblast cells HL-60s (ATTC, CCL-240) were cultured in differentiation media consisting of Roswell Park Memorial Institute (RPMI) 1640 or Iscove’s Modified Dulbecco’s medium (IMDM) supplemented with 20% fetal bovine serum (FBS, Cytiva HyClone), 2 mM L-glutamine (Corning), 100 U ml−1 penicillin, and 100 μg ml−1 streptomycin, 1.25% dimethyl sulfoxide (DMSO), and 1 μM All-trans retinoic acid (ATRA) for 3 days. HL-60 differentiation into NLCs using DMSO and ATRA has been well established and widely used in numerous studies to model neutrophil function in vitro (Abi Abdallah et al., 2012; Bliss et al., 2001; Tasseff et al., 2017). Confirmation of HL-60 differentiation into neutrophil-like cells (NLCs) was assessed by flow cytometry using a human anti-CD11b antibody (BioLegend, ICRF44).

### *T. gondii* culture and neutrophil infection

Human foreskin fibroblasts (HFFs) were cultured in Dulbecco’s Modified Eagle Medium supplemented with 2 mM L-glutamine (Corning), 100 U ml−1 penicillin, 100 μg ml−1 streptomycin (HyClone), and 10% FBS. Cells were maintained at 37°C with 5% CO_2_. Type I (RH) and type II (PA7) *T. gondii* tachyzoites expressing green fluorescent protein (GFP) were maintained by serial passage in HFF monolayers as previously described (Gov et al., 2017; Lima et al., 2018; Morgado et al., 2011). Parasites were harvested from HFFs by syringe lysis (27-gauge needle) followed by filtration through a 0.5 μm filter to remove host cell debris. NLCs were infected at a multiplicity of infection (MOI) of 2, while mock infections used an equal volume of media without parasites. To ensure comparable infection efficiency between *T. gondii* strains, flow cytometry was performed to quantify intracellular GFP signal.

### Sytox assay

NLCs (2.0 × 10^5^) were incubated for 15 minutes in the presence of 5 µM Sytox Orange (Invitrogen) before infection. Experiments used 4% FBS (Hyclone) or 4% human serum (HS, MP Biomedical). Cells were infected with type I or type II *T. gondii* tachyzoites at an MOI of 2, whereas mock-infected cells were incubated with RPMI 1640 only. As a positive control, NLCs were treated with 100 ng PMA. Fluorescence was measured every hour for 8 hours using a Molecular Devices SpectraMax iD3 plate reader at 570 nm. Fluorescence values were subtracted from the background fluorescence of a blank, and the area under the curve was calculated for each experiment.

### Immunofluorescence microscopy

NLCs were placed in wells containing poly-L-lysine-coated coverslips (Corning) at a concentration of 4 × 10^5^/mL and infected with type I or type II *T. gondii* parasites at an MOI of 2 with either FBS or HS. As a positive control, NLCs were treated with 100 ng PMA. Coverslips were harvested, fixed in 4% paraformaldehyde (PFA) for 30 minutes, and permeabilized with 0.5% Triton X-100. Blocking was performed in 5% goat serum, and the samples were stained with a primary antibody against citrullinated histone 3 (Novus Biological Sciences, ICFR44, 1:1000 dilution) overnight. Afterward, cells were stained with a secondary antibody, anti-rabbit IgG-AF594 (5 µg/mL) for 2 hours and mounted with antifade DAPI mounting medium. Coverslips were imaged on a Nikon Eclipse Ti Confocal Microscope. Images were taken using the NIS-Elements software at 40X, 60X, and 100X. Images were then processed in ImageJ (Fiji), where z-stack images were compiled into a mean fluorescence image. Signal quantification was performed utilizing ImageJ, and signal intensity was normalized to mock-infected controls in Prism GraphPad.

### Cell viability and total cell counts

To assess cell viability and total cell numbers, NLCs were collected and washed once in phosphate-buffered saline (PBS). Cells were then resuspended in a 1:100 dilution of Zombie Violet fixable viability dye (BioLegend) prepared in PBS and stained for 20–30 minutes on ice, protected from light. Cells were subsequently fixed in 4% PFA for downstream analysis. Total cell counts and viability were quantified by flow cytometry, using Zombie Violet fluorescence to distinguish live from dead cells. As a positive control, NLCs were treated with 100 ng PMA and processed at the respective time point.

### ATP release assay

A total of 2 ×10^4^ NLCs were plated on a 96-well plate and infected with type I or type II *T. gondii* tachyzoites in the presence of HS, with extracellular ATP measurements taken every hour for 6 hours. As a positive control, NLCs were treated with 100 ng PMA. The culture supernatant was assessed for extracellular ATP release using a commercial kit (Promega) according to the manufacturer’s instructions. Luminescence values were plotted in GraphPad Prism at 1-hour intervals over 6 hours. The area under the curve was calculated for each experiment.

### Blocking of complement receptor 3 (CR3)

NLCs were incubated with an anti-CD11b blocking antibody (BioLegend, Ultra-Leaf ICRF44) at 40 μg/mL for 1 hour to block CD11b/CR3 prior to infection. Cells were then used in Sytox or immunofluorescent microscopy experiments. Confirmation of CR3 blocking by epitope occupancy was assessed by flow cytometry using a human anti-CD11b antibody (BioLegend, ICRF44).

### Statistics

Statistical analysis was performed using GraphPad Prism and a minimum of 4 independent experiments. Values were expressed as means, and error bars reflect standard deviation. Analysis of variance (ANOVA), followed by Bonferroni’s test, was used to compare means. Differences were considered significant when *P* < 0.05.

## Results

### Neutrophil-like cells serve as a model to study neutrophil-mediated responses to *T. gondii* infection

*T. gondii* infection induces NLCs to release neutrophil extracellular traps (NETs) (Abi Abdallah et al., 2012; Bliss et al., 2001). Thus, to investigate the mechanisms associated with this process, the promyeloblast cell line HL-60 was differentiated into neutrophil-like cells (NLCs) by treatment with DMSO and retinoic acid for 3 days, as previously described (Bergersen et al., 2023; Miranda et al., 2021). HL-60s are promyeloblasts that express little to no CD11b before maturation into NLCs, and the differentiation protocol was confirmed by the resulting 93% CD11b expression and over 95% viable cells (Fig. 1A). Additionally, DAPI staining was used to confirm nuclear morphological changes during differentiation. Undifferentiated HL-60s presented a large, round nucleus, while differentiated NLCs gained the characteristic multilobed nucleus of neutrophils (Fig. 1B). NLCs were infected with green fluorescent protein (GFP)-expressing *T. gondii* (type I or type II), and the infection efficiency and cell viability were assessed over time. These conditions resulted in an infection efficiency of approximately 42% at 4 hours post-infection, with a viable cell percentage of approximately 93% (Fig. 1C).

**Figure 1.**
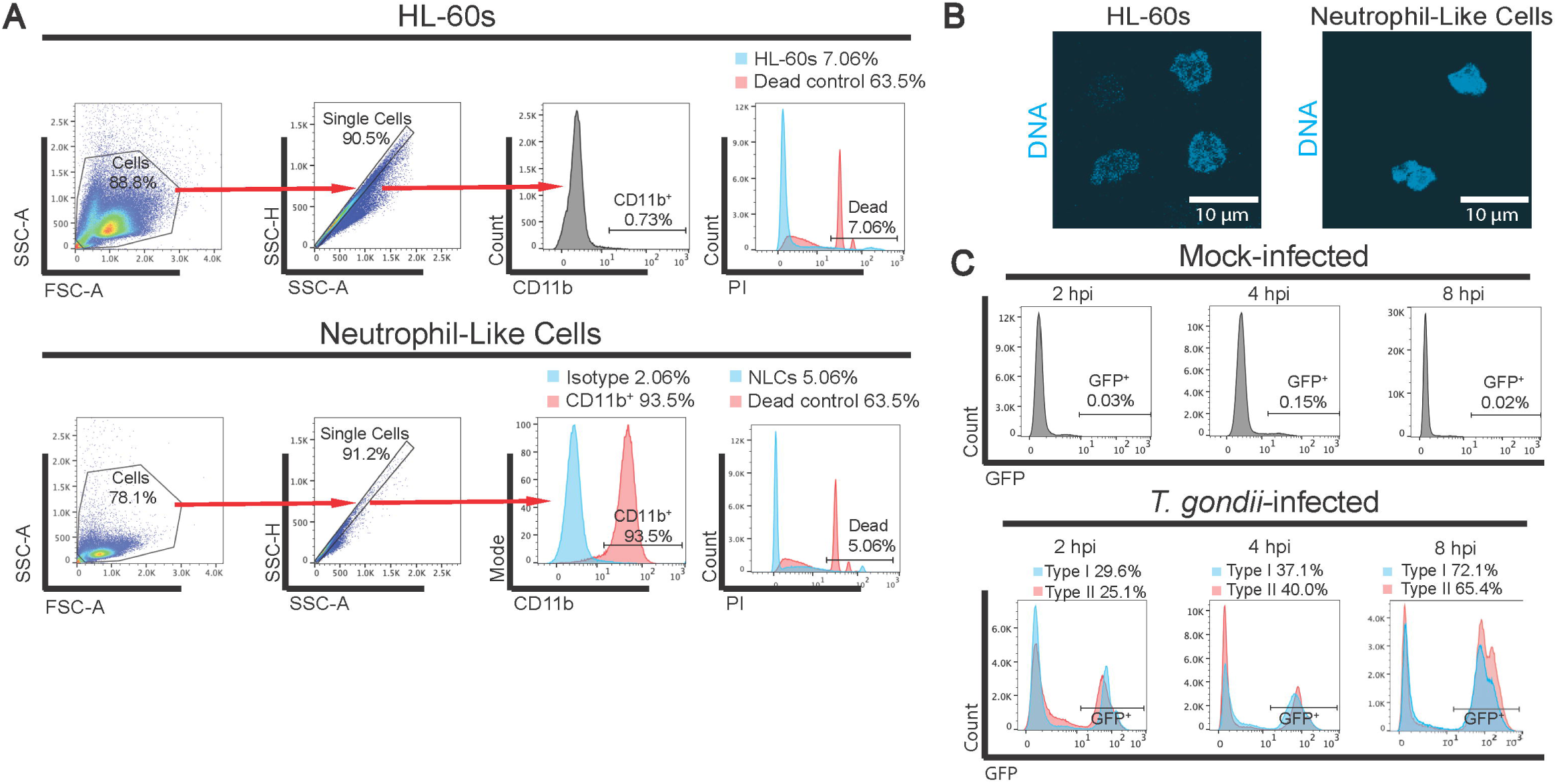
Differentiation of HL-60 cells into neutrophil-like cells (NLCs). (A) HL-60 cells were differentiated into NLCs by treatment with DMSO and retinoic acid for 3 days. Flow cytometry analysis was used to confirm CD11b expression and cell viability (PI^-^ cells). (B) Nuclear morphology was confirmed by staining HL-60 and NLC cells with DAPI, followed by confocal microscopy analysis at 100X (C) Infection efficiency was analyzed by flow cytometry in response to type I and type II *T. gondii* at 2, 4, and 8 hours post-infection (hpi). (A and C) These controls were performed for all experiments. (B) Representative images from 3 independent experiments are shown.

### *T. gondii*-infected NLCs release NETs

We next compared the ability of NLCs to release NETs in response to type I or type II *T. gondii* infection. NLCs were infected at an MOI of 2 and cultured in the presence of Sytox, an impermeable fluorescent dye that binds extracellular DNA. Fluorescence readings taken hourly for 8 hours showed an increase in extracellular DNA in response to type I *T. gondii* and PMA, a well-characterized inducer of NETosis (Fig. 2A and 2B). Although extracellular DNA increased, its presence could result from NETosis or merely from cell death. Additionally, a notable limitation was the inability to detect extracellular DNA following type II *T. gondii* infection, potentially due to differences in the magnitude or kinetics of the NET response elicited by type II parasites. Thus, we next confirmed the presence of NETs by performing confocal immunofluorescence microscopy for DNA and citrullinated histone 3 (H3Cit), a key marker of NETosis. NET structures were observed in response to both type I and type II *T. gondii* at 4 and 16 hours post-infection (Fig. 2C). Additionally, PMA also induced NET structures at both time points, but the presentation of NETs was less web-like and more consistent with the characterized suicidal NETosis of cell swelling and bursting (Fig. 2C). Interestingly, *T. gondii* infection did not cause NLC nuclei to swell, a characteristic of suicidal NETosis before the cell membrane bursts and leads to cell death, thus confocal imaging suggested that NLC nuclei remain intact even after NET release.

**Figure 2.**
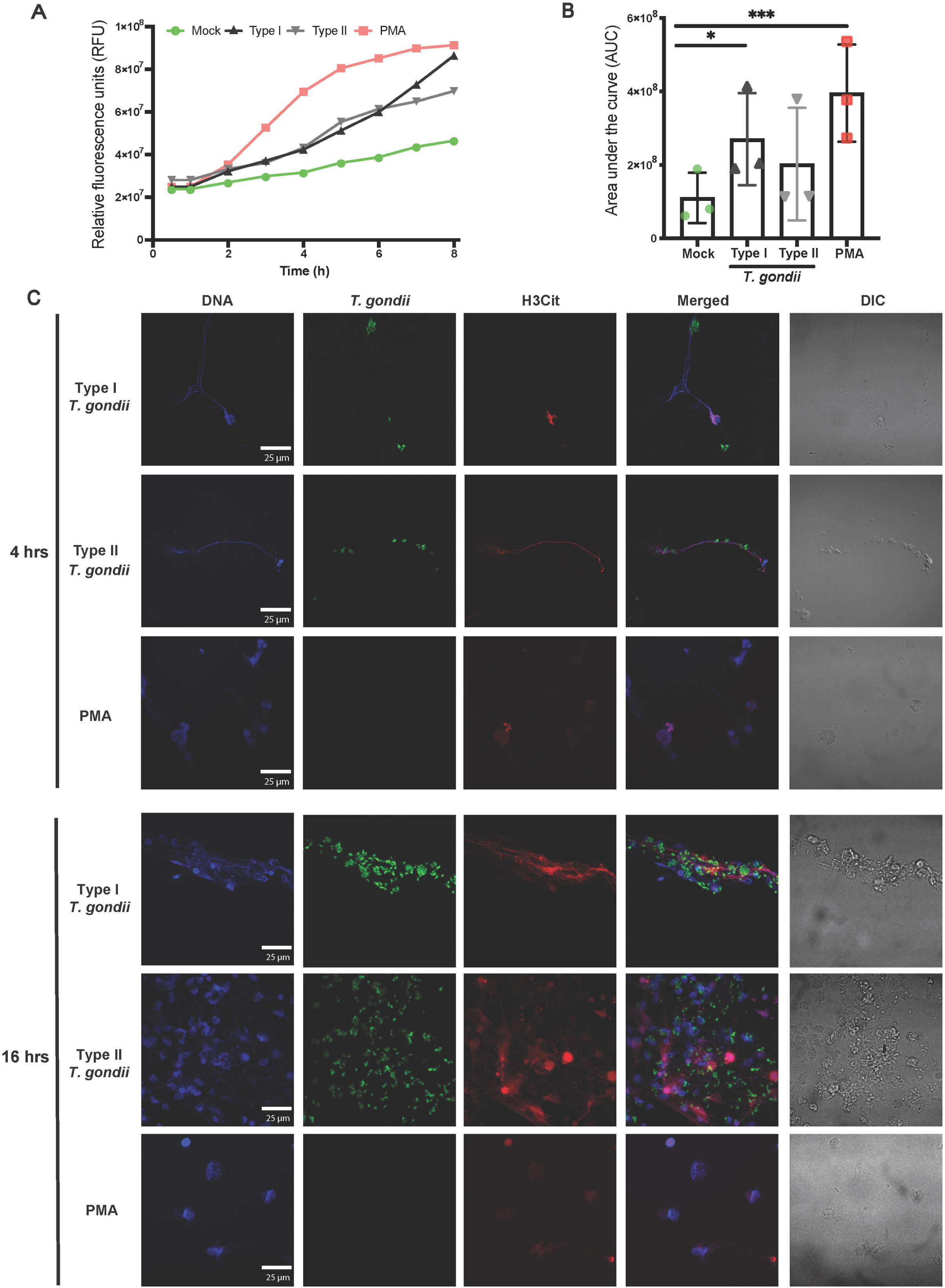
Type I and type II *T. gondii* induce neutrophil extracellular traps (NETs) release by neutrophil-like cells (NLCs). NLCs were either mock-infected, infected with GFP-expressing *T. gondii* (type I RH or type II PA7), or treated with PMA. (A) Extracellular DNA was measured for 8 hours in NLCs infected with *T. gondii* using SYTOX dye, and PMA-stimulated NLCs served as a positive control for suicidal NETosis. (B) Area under the curve (AUC) quantification. (C) NET formation was confirmed by immunofluorescence microscopy using an antibody against citrullinated histone 3 (H3Cit) and DAPI staining in *T. gondii*-infected or PMA-treated NLCs at 4 and 16 hours. Values are expressed as means ± standard deviation. *, P<0.05, *** P<0.001 (one-way ANOVA followed by Bonferroni’s post hoc test in panel B). (A and C) Representative curves and images from 3 independent experiments, respectively. (B) Experiments were performed 3 independent times, and the combined data are shown.

Together, these findings indicate that NLCs can serve as a model for investigating molecular mechanisms associated with the NETosis immune response against type I and type II *T. gondii*. Additionally, the observation of intact nuclei at 16 hours prompted an investigation into whether vital NETosis is the primary pathway used by NLCs in response to *T. gondii*.

### Vital NETosis is induced by *T. gondii*

Suicidal NETosis is well known to disrupt neutrophil plasma membranes, leading to cell death (Remijsen et al., 2011; Takei et al., 1996; Yipp & Kubes, 2013). With plasma membrane integrity being one of the most significant differences between suicidal NETosis and vital NETosis, we examined the plasma membrane integrity of *T. gondii*-infected NLCs. Cells were infected for 2, 4, 6, 8, or 16 hours, followed by staining with the viability dye Zombie Violet and flow cytometry analysis to assess viability by comparing Zombie Violet-positive cell percentages. Samples were compared to a heat-killed positive control (Fig. 3A). NLCs infected with type I and type II *T. gondii* showed approximately 2.4% and 1.9% dead cells, respectively, both of which are comparable to the 4.8% dead cells in mock-infected samples, and heavily contrasted with PMA, which showed 16.9% dead cells at 2 hours, thus suggesting that *T. gondii* and PMA trigger different NETosis pathways (Figs. 3A and 3B). At 16 hours post-infection, we found similar results with dead cell percentages averaging 7.5% for mock-infected, 7.4 % and 7.6% for type I and type II *T. gondii*-infected NLCs, respectively, but heavily contrasted to PMA, which averaged 64% (Fig. 3B). To ensure that the percentages of dead cells were not skewed by cellular losses during centrifugations, total cell counts were obtained, showing similar counts at each time point across all *T. gondii*-infected samples, as compared to PMA samples wherein cell numbers decreased due to cell death (Fig. 3C). To further assess plasma membrane integrity, extracellular ATP activity was quantified by luminescence readings taken every hour for 6 hours and compared to those of digitonin, a powerful detergent, and PMA. Extracellular ATP increased dramatically in PMA and digitonin samples, whereas in type I *T. gondii*-infected NLCs, ATP levels were marginally elevated, possibly due to ATP released by activated NLCs as they responded to the infection (Fig. 3D and E).

**Figure 3.**
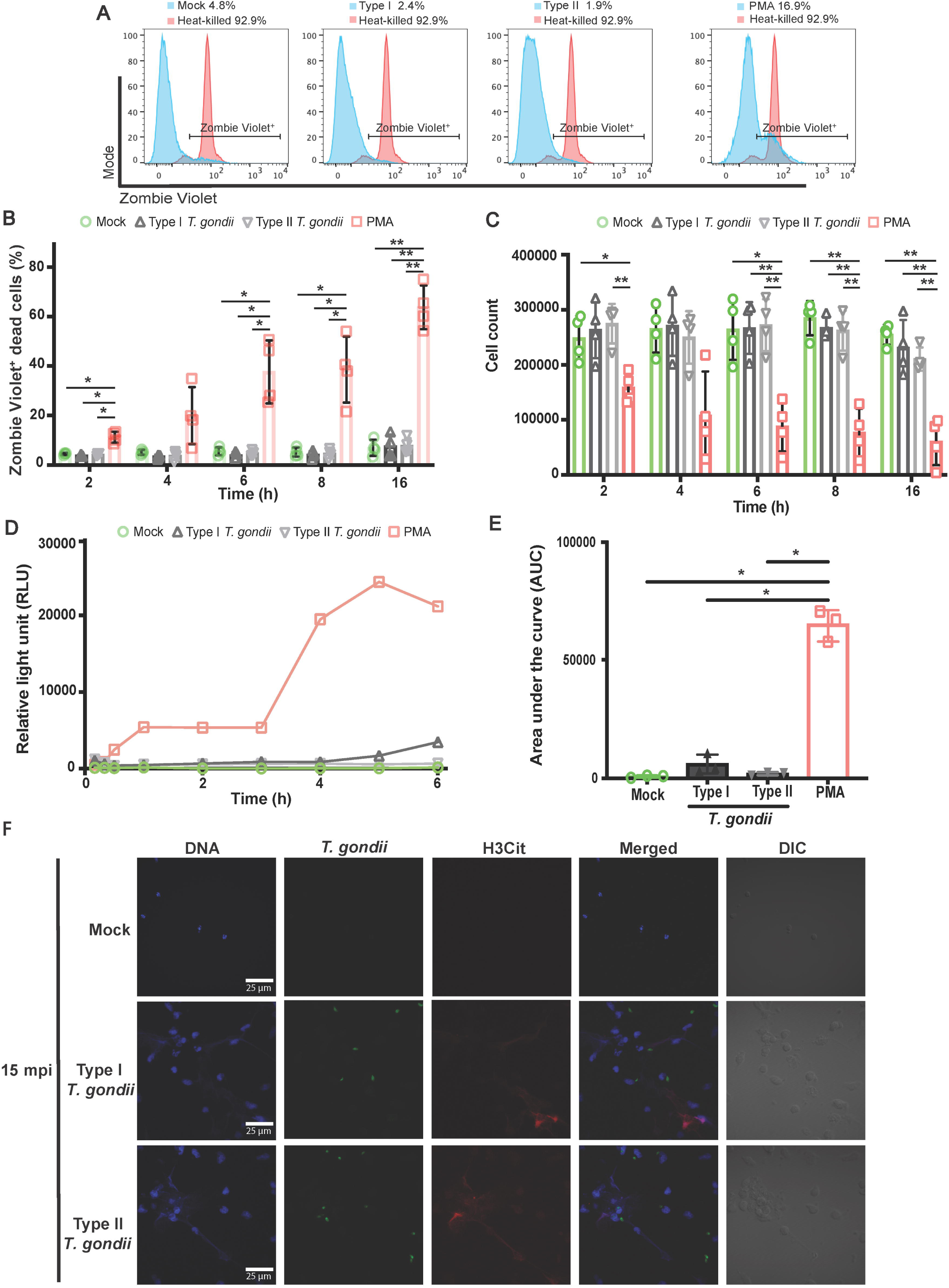
***T. gondii*-infected neutrophil-like cells (NLCs) maintain membrane integrity and release neutrophil extracellular traps (NETs) through vital NETosis.** NLCs were either mock-infected, infected with GFP-expressing *T. gondii* (type I RH or type II PA7), or treated with PMA. (A) Plasma membrane integrity was assessed by flow cytometry using Zombie Violet dye for 16 hours. (B) Percentage of dead NLCs over time. (C) Total cell counts over time. (D) Extracellular ATP levels were measured for 6 hours in the culture. (E) Area under the curve (AUC) quantification. (F) Vital NETosis was further confirmed by immunofluorescence microscopy at 15 minutes post-infection (mpi). Values are expressed as means ± standard deviation. *, P<0.05, ** P<0.01 (two-way ANOVA, followed by Tukey’s post hoc test in panels B, C, and E). (A, D and F) Representative flow cytometry, time course, and images from 3-4 independent experiments are shown. (B, C, and E) Experiments were performed 3-4 independent times, and the combined data are shown.

Suicidal NETosis occurs, on average, 2 hours after stimulation to induce NET release, whereas vital NETosis has been documented to occur quickly, within minutes after stimulation (Pilsczek et al., 2010a) To confirm that *T. gondii* tachyzoites induce vital NETosis, we performed immunofluorescence microscopy at early time points (15 and 30 minutes). NET structures entrapping *T. gondii*, composed of DNA and H3Cit, were observed as early as 15 minutes post-infection in the extracellular space (Fig. 3F). Together, these data indicate that NETs are released against *T. gondii* without disrupting the plasma membrane, thus indicating activation of the vital NETosis pathway.

### Complement receptor 3 (CR3) mediates *T. gondii*-induced vital NETosis

Vital NETosis has been implicated in responses to bacterial pathogens following complement receptor 3 (CR3) activation (Palmer et al., 2016a) but the role of this receptor in responses to *T. gondii* remains unknown. Given that complement opsonization enhances CR3-mediated NETosis in other infection models, we hypothesized that serum opsonins may similarly increase NET release during *T. gondii* infection. To evaluate opsonins’ role in mediating neutrophil responses to *T. gondii*, we assessed NETosis using the Sytox extracellular DNA assay. NLCs infected in the presence of human serum released more extracellular DNA than those incubated with FBS, suggesting that opsonins enhance NET formation in response to *T. gondii* (Fig. 4A and B).

**Figure 4.**
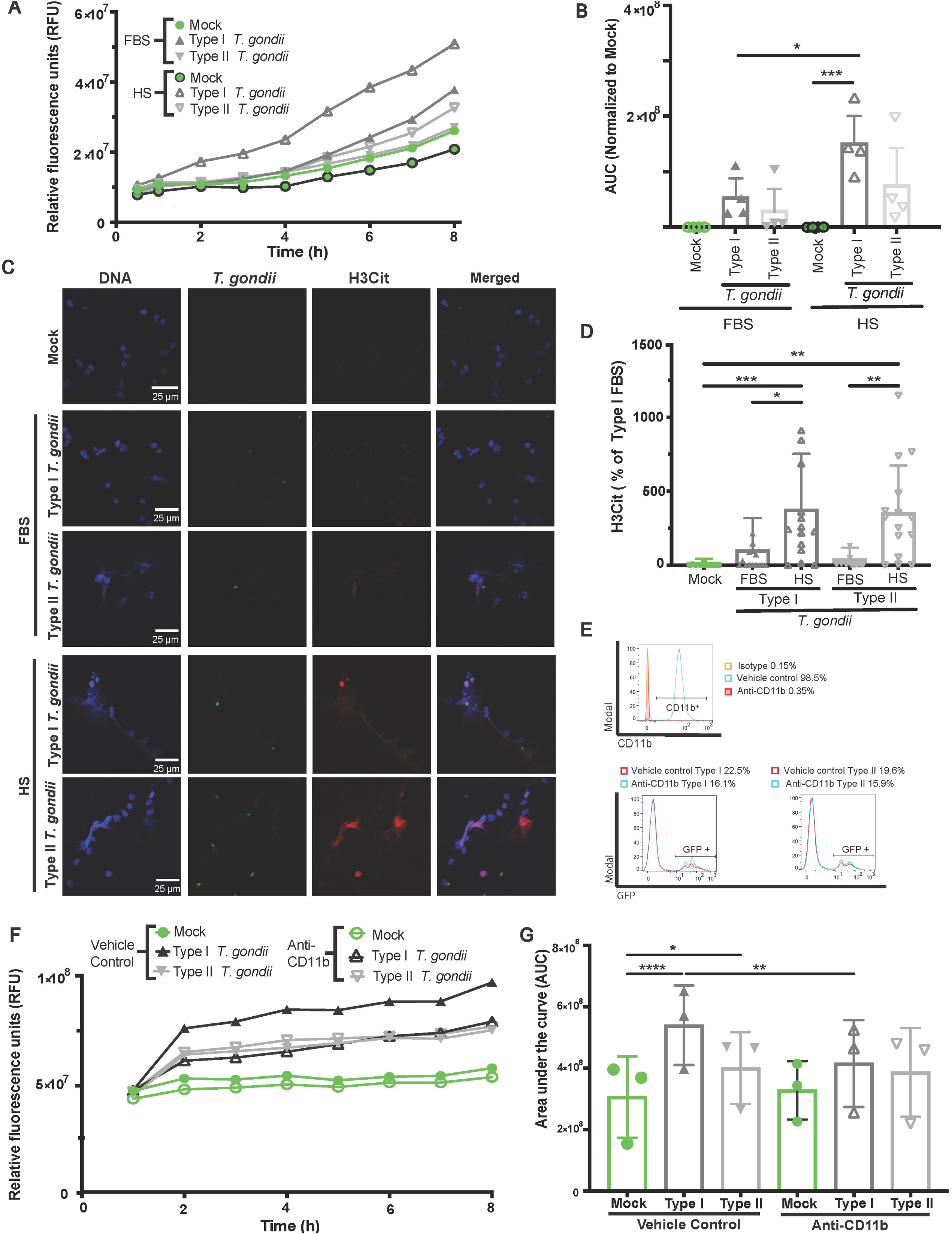
Complement receptor 3 mediates opsonin-dependent NET release in response to *T. gondii*. NLCs were either mock-infected or infected with GFP-expressing *T. gondii* (type I RH or type II PA7) in the presence of fetal bovine serum (FBS) or human serum (HS). (A) Extracellular DNA was measured for 8 hours. (B) Area under the curve (AUC) quantification. (C) Immunofluorescence microscopy using an antibody against citrullinated histone 3 (H3Cit) and DAPI staining at 45 minutes post-infection (mpi). (D) Quantification of relative H3Cit levels. (E) Flow cytometry confirmation of CR3 blocking and infection efficiency in the presence of blocking antibodies. (F) Extracellular DNA was measured for 8 hours in *T. gondii*-infected NLCs previously treated with anti-CD11b blocking antibodies or vehicle control (DMSO). (G) Area under the curve (AUC) quantification. Values are expressed as means ± standard deviation. *, P<0.05, ** P<0.01, *** P<0.001, **** P<0.0001 (two-way ANOVA, followed by Tukey’s post hoc test in panels B, D, and G). (A, C, E, and F) Representative time courses, images, and flow cytometry from 3-4 independent experiments are shown. (B and G) Experiments were performed 3-4 independent times, and the combined data are shown. (D) A total of 15 images per condition from 3 independent times were analyzed.

When comparing immunofluorescence microscopy images taken at 45 minutes post-infection of type I- and type II *T. gondii*-infected NLCs cultured in FBS or HS, we observed greater NET release under HS conditions (Fig. 4C). A relative quantification showed a notable increase in H3Cit levels compared to mock controls, indicating once again NET formation in response to infection. Specifically, HS-cultured NLCs infected with type I or type II *T. gondii* displayed higher H3Cit levels compared to FBS-cultured NLCs (Fig. 4D), indicating that active serum components in HS may facilitate or amplify the NETosis response.

To determine whether CR3 activation contributed to this enhanced response, neutrophils were treated with CR3-blocking antibodies (anti-CD11b) prior to infection. Epitope occupancy and comparable infection efficiency in the presence of anti-CD11b were confirmed by flow cytometry (Fig. 4E). Blocking CR3 significantly reduced extracellular DNA release compared with vehicle-treated cells (Fig. 4F and G), demonstrating that CR3 plays a critical role in *T. gondii* crosslinking and opsonin-mediated enhancement of NETosis against the parasite.

## Discussion

Investigating the mechanisms by which NETs are produced against *T. gondii* is grounded in the need to understand how innate immune cells respond to this parasite. This foundational knowledge can inform the development of therapeutics that specifically enhance immune function in infected cells, helping control *T. gondii* spread and attenuate disease pathogenesis. The production of extracellular traps by neutrophils was demonstrated years ago (Brinkmann et al., 2004); however, the exact mechanisms underlying this response have not been fully elucidated. Here, we showed that *T. gondii* induces NET formation through the CR3-mediated vital NETosis pathway. Using HL-60–derived NLCs as an in vitro model, we showed that both type I (RH) and type II (PA7) *T. gondii* tachyzoites stimulated extracellular DNA release. Using immunofluorescence microscopy, we confirmed the presence of NET structures marked by H3Cit and DNA produced against the parasite. Critically, NET-releasing NLCs retained plasma membrane integrity and viability, distinguishing this response from suicidal NETosis and implicating vital NETosis as the operative pathway. Although previous studies suggested that type I *T. gondii*-infected primary neutrophils release both classical and rapid NETs (Macedo et al., 2023) these studies did not investigate whether the integrity of the cellular plasma membrane or nuclei remained intact. Our study furthers this investigation by confirming plasma membrane integrity and cell viability during *T. gondii-*induced NETosis, in addition to investigating type II *T. gondii* NET release.

Differentiating HL-60 promyeloblasts into NLCs with DMSO and retinoic acid was confirmed by elevated CD11b expression and multilobed nuclei, consistent with previous studies showing that retinoic acid improves both differentiation and NET formation (Tasseff et al., 2017). The infection rate at 4 hours post-infection, along with sustained cell viability, established a clear baseline for studying NETosis pathways without interference from significant cell death due to the lytic nature of *T. gondii*.

Sytox-based quantification of extracellular DNA showed a time-dependent increase in DNA released from NLCs exposed to both *T. gondii* strains, although type I produced more NETs than type II, mirroring the response in PMA-stimulated positive controls. However, immunofluorescence microscopy revealed a qualitative difference: *T. gondii*-infected NLCs did not exhibit nuclear swelling that typically precedes membrane rupture in suicidal NETosis, and their nuclei remained intact even after NET release at 16 hours post-infection. This morphological finding aligns with our viability assay, as Zombie Violet staining indicated a cell death rate in *T. gondii*-infected NLCs comparable to mock-infected NLCs and all three groups significantly differing from PMA-treated NLCs. Additionally, extracellular ATP measurements supported membrane integrity, with only minor ATP release from *T. gondii*-infected NLCs compared to the significant release caused by PMA. Altogether, these membrane integrity assessments support the conclusion that *T. gondii* induces vital NETosis in NLCs.

The detection of NET structures at 15 and 30 minutes post-infection is particularly noteworthy. PMA suicidal NETosis formation takes at least 2 hours to occur when PAD4-mediated histone citrullination, chromatin decondensation, and plasma membrane rupture (Clark et al., 2018; Kubes, 2018; Yipp & Kubes, 2013). Since NET release within 15–30 minutes cannot be explained by the suicidal pathway, it instead supports the vesicular extrusion mechanism associated with vital NETosis (Brinkmann et al., 2004; Pilsczek et al., 2010a). In this process, nuclear envelope budding results in DNA-filled vesicles that are released and rupture outside the cell, while the neutrophil remains intact and functional. This timing is aligned with previous studies of vital NETosis triggered by *Staphylococcus aureus* and platelets, and our work broadens those findings to include a protozoan parasite (Brinkmann et al., 2004; Pilsczek et al., 2010b).

The enhanced release of extracellular DNA observed with HS, compared with FBS, suggests that human-specific opsonins, such as complement proteins and immunoglobulins, contribute to *T. gondii* crosslinking and neutrophil responses. Inhibition of CR3 (CD11b/CD18) with anti-CD11b antibodies significantly reduced this serum-dependent increase, highlighting CR3 as an important receptor involved in *T. gondii*-induced vital NETosis. CR3 is a versatile integrin that binds iC3b/C3b-opsonized targets and is linked to neutrophil adhesion, phagocytosis, and oxidative burst (Chen et al., 2021; Clark et al., 2018). A few studies have demonstrated the role of CR3 in mediating vital NETosis (Chen et al., 2021; Palmer et al., 2016b; Pilsczek et al., 2010b; Yipp & Kubes, 2013) and our findings now identify vital NETosis in response to *T. gondii* infections as another important downstream effect. The complement-CR3 interaction could be especially important in vivo during acute intestinal infections, when complement activation is rapid, and neutrophils are among the initial immune responders (Sikorski et al., 2021).

A key aspect of vital NETosis is that surviving neutrophils retain their effector functions, such as chemotaxis, phagocytosis, and cytokine secretion. These functions are lost during suicidal NETosis. If *T. gondii*-stimulated neutrophils in vivo follow the vital NETosis pathway described here, they could both release NETs to trap extracellular tachyzoites and continue to produce IL-12 and other cytokines that support the IFN-γ–dependent adaptive immune response (Biswas et al., 2017; Miranda et al., 2021). This response is essential for disease control and for conversion to bradyzoites as the infection transitions into the chronic stage (Bergersen et al., 2023; Biswas et al., 2017; Kim et al., 2007; Miranda et al., 2021). Consistent with this model, studies have shown that *T. gondii*-induced NETs do more than physically restrain the parasite: they also amplify the innate inflammatory response and help prime the adaptive response (Biswas et al., 2017; Miranda et al., 2021). This supports the idea that NET release by neutrophils that remain functional could contribute to the type 1 immunity needed to control infection, rather than acting solely as an antiparasitic trapping mechanism. Such dual roles may explain why neutrophil depletion leads to increased parasite burdens and weakened type 1 cytokine responses in mouse models, effects that cannot be solely attributed to NETs (Bliss, 1999; Bliss et al., 2001).

While the use of NLCs is a well-established neutrophil model (Bergersen et al., 2023; Brinkmann et al., 2004; Burn et al., 2021; Tasseff et al., 2017) the use of this cell line may not fully recapitulate how primary human neutrophils respond to *T. gondii* infections. Previous studies have demonstrated that *T. gondii* infection of primary human neutrophils delays apoptosis and prolongs cell viability, while these cells still undergo NETosis (Lima et al., 2021), thus suggesting that the mechanisms described in this study are also present in primary neutrophils. Future investigations should explore the potential roles of additional complement receptors, such as CR1 and CR4, as well as Fc receptors, which may act in concert with or independently of CR3 to facilitate NET release in response to *T. gondii* infection, mechanisms that, while uncharacterized in *T. gondii,* have been documented in other pathogens (Alemán et al., 2016; Bennouna et al., 2003; Chen et al., 2021). In summary, this study demonstrates that *T. gondii* triggers CR3-mediated vital NETosis in NLCs. These results contribute to expanding our understanding of how neutrophils might contribute to the initial immune response to toxoplasmosis and suggest new directions for exploring their varied roles during parasitic infections.

## Acknowledgments

We would like to thank Dr. Melissa Lodoen from the University of California, Irvine, for the gift of RH and PA7 *T. gondii* parasites. We also thank all members of the Lima and Mercer labs at Cal Poly Pomona for helpful discussions on this project.

This work was supported by the National Institutes of Health (NIAID grant R16AI184345 to Tatiane S. Lima and NIGMS grant T32GM137812 to Marco A. Martinez). The funders had no role in study design, data collection, and interpretation, or the decision to submit the work for publication.

